# Permanent deconstruction of intracellular primary cilia in differentiating granule cell neurons

**DOI:** 10.1101/2023.12.07.565988

**Authors:** Carolyn M. Ott, Sandii Constable, Tri M. Nguyen, Kevin White, Wei-Chung Allen Lee, Jennifer Lippincott-Schwartz, Saikat Mukhopadhyay

## Abstract

Primary cilia on granule cell neuron progenitors in the developing cerebellum detect sonic hedgehog to facilitate proliferation. Following differentiation, cerebellar granule cells become the most abundant neuronal cell type in the brain. While essential during early developmental stages, the fate of granule cell cilia is unknown. Here, we provide nanoscopic resolution of ciliary dynamics *in situ* by studying developmental changes in granule cell cilia using large-scale electron microscopy volumes and immunostaining of mouse cerebella. We found that many granule cell primary cilia were intracellular and concealed from the external environment. Cilia were disassembed in differentiating granule cell neurons in a process we call cilia deconstruction that was distinct from pre-mitotic cilia resorption in proliferating progenitors. In differentiating granule cells, ciliary loss involved unique disassembly intermediates, and, as maturation progressed, mother centriolar docking at the plasma membrane. Cilia did not reform from the docked centrioles, rather, in adult mice granule cell neurons remained unciliated. Many neurons in other brain regions require cilia to regulate function and connectivity. In contrast, our results show that granule cell progenitors had concealed cilia that underwent deconstruction potentially to prevent mitogenic hedgehog responsiveness. The ciliary deconstruction mechanism we describe could be paradigmatic of cilia removal during differentiation in other tissues.

## INTRODUCTION

A singular, specialized, signal detecting organelle, called the primary cilium, protrudes from the surface of most cells including neurons (Green and Mykytyn, 2014; Louvi and Grove, 2011; Ott et al., 2023; Rosenbaum and Witman, 2002; Wu et al., 2023). Receptors and effectors compartmentalized in the cilium mediate efficient detection and transduction of external signals (Anvarian et al., 2019; Nachury and Mick, 2019). Even deep in tissues, cilia perceive and respond to signals that can initiate developmental programs or alter cell activity. Despite the significant functions of cilia, variations in cilia structure within tissues have just begun to be studied (Mill et al., 2023). In the brain and spinal cord, for example, cilia abundance and length can change (Das and Storey, 2014; Tu et al., 2023). Because cilia sense growth factors, neuropeptides and neuromodulators, changes in cilia can impact neurodevelopment, neuronal function, neuronal circuit connectivity, neuronal excitability, and neuropathology (Bowie and Goetz, 2020; Guo et al., 2017; Higginbotham et al., 2012; Kumamoto et al., 2012; Lee and Gleeson, 2011; Sheu et al., 2022; Suciu and Caspary, 2021; Tereshko et al., 2021; Wilsch-Brauninger and Huttner, 2021; Youn and Han, 2018). To understand these processes, we need to understand how changes in cilia ultrastructure contribute to neurodevelopmental programs, especially in intact tissue.

Approximately eighty percent of adult human brain neurons are packed into the inner layer of the cerebellum (Herculano-Houzel, 2009). These neurons are a singular type of neuron, called granule cells (GCs). GC neurons are particularly interesting from a cilia perspective because their expansion from immature progenitors occurs in response to sonic hedgehog (SHH), a soluble ligand detected by receptors in the ciliary membrane, which triggers progenitor proliferation (Dahmane and Ruiz i Altaba, 1999; Wallace, 1999; Wechsler-Reya and Scott, 1999). Mutations that disrupt ciliogenesis lead to cerebellar hypoplasia and abnormalities in foliation (Chizhikov et al., 2007; Spassky et al., 2008). Upon onset of differentiation GCs stop responding to SHH, begin to extend axons, and migrate along glial processes toward Purkinje neuron cell bodies. At this early stage of differentiation, reductions in cilia length and frequency have been reported (Chang et al., 2019; Di Pietro et al., 2017; Ong et al., 2020), raising the possibility that modulation of ciliation could be involved in the differentiation of these cells. However, the magnitude and duration of cilia modulation during differentiation have not yet been determined. Addressing this question with detailed ultrastructural analysis has been challenging because cilia and centrosomes are small, singular structures.

To characterize morphological changes in cilia within the cerebellum during neurogenesis we mapped and quantified centrosomes and cilia in hundreds of GCs using large volume electron microscopy (EM) datasets of adult and developing mouse cerebella (Nguyen et al., 2023; Wilson et al., 2019). By combining high resolution EM with the molecular specificity of immunofluorescence imaging, we quantified modulation in cilia length and abundance, obtaining unprecedented nanoscopic images of centrosomes and cilia across the span of neurogenesis. Our findings reveal a process for cilia loss during GC differentiation that we call cilia deconstruction. We report surprising intermediates and end states in the ciliary deconstruction process. In mature GCs centriole distal appendages were anchored to the plasma membrane without forming cilia. Together, the results demonstrate for the first time dynamic cilia architecture at the nanoscale across a tissue and offer insights into cilia maintenance and modulation pathways likely relevant to other tissues during development.

## RESULTS

### Primary cilia length and frequency are decreased during GC neurogenesis

To investigate cilia status throughout the entire progression of GC differentiation in the postnatal developing cerebellum, we immunostained and imaged cerebellar sections from postnatal day 9 (P9) mice using an antibody to ARL13B (Figure 1A). ARL13B is an atypical GTPase that is highly enriched in cilia (Caspary et al., 2007; Duldulao et al., 2009). We stained for SOX9 to identify glia (Farmer et al., 2016; Sun et al., 2017; Vong et al., 2015), and the cell cycle inhibitor P27^KIP1^ to delineate differentiating and mature GCs from cycling GC progenitors (Miyazawa et al., 2000). The progression of GC neuronal maturation coincides with migration deeper into the tissue. Therefore, in a single sagittal section the maturation state of each P27^KIP1^ expressing GC can be inferred by the depth of the GC in the cerebellum (Leto et al., 2016). As expected, GC progenitors that populated the external granule layer (EGL) of the emerging cerebellum lacked P27 ^KIP1^ expression, while GCs newly committed to differentiation that expressed P27^KIP1^ had exited the proliferation zone and were located in the inner EGL (Figure 1A). Differentiating GCs migrating through the molecular layer (ML), past the Purkinje soma in the Purkinje cell layer (PCL), were also expressing P27^KIP1^, as were GC neurons that were located in the internal granular layer (IGL) in the latest stages of neurogenesis and maturation (Figure 1A).

**Figure 1.**
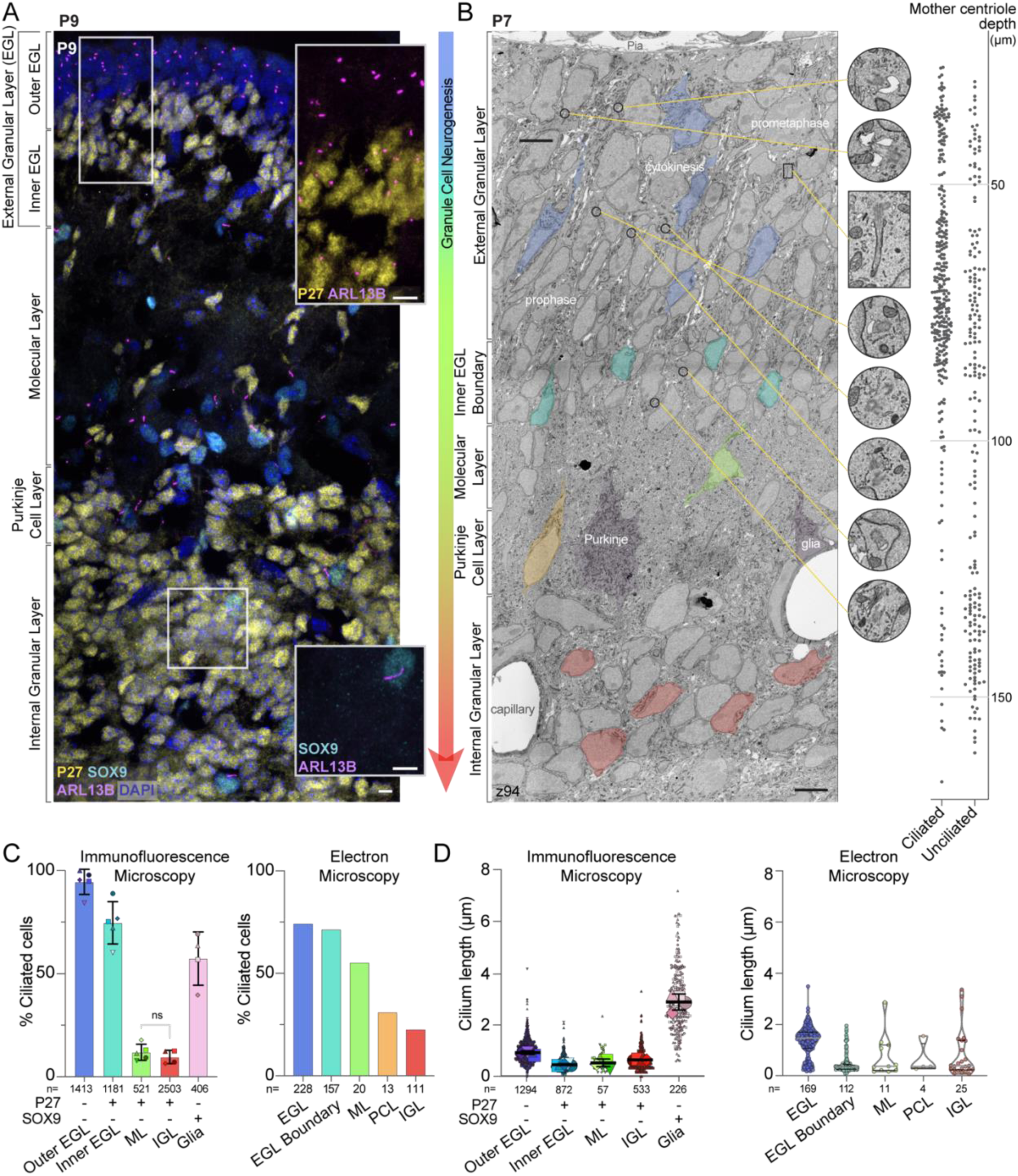
Differentiating GCs initially have short cilia that are lost as neurons mature. Gradual (A) A sagittal section of P9 mouse cerebellum immunostained with antibodies to the GC differentiation marker P27^KIP^ (yellow), the cilia marker ARL13B (magenta), the glial marker SOX9 (cyan), and counterstained with DAPI (blue) then imaged with spinning disc confocal microscopy. The layers of the developing cerebellum are indicated on the left. Scale bar: 10 μm. (B) A single slice of the P7 large serial-section scanning EM volume with GCs in different stages of differentiation are highlighted. The phase of mitotic cells in the EGL are superimposed on the GCs. In addition, cropped images of centrosomes and cilia from the panel are magnified and the location of each image is indicated by a yellow line. On the right side of the image, the depth of each ciliated and unciliated mother centriole is plotted. Scale bar: 5 μm; diameter of zoom regions: 1.6 μm. (C) Cilia frequency is quantified from measurements of widefield immunofluorescent images (left; 3 sections from each of 4 or 5 animals) and from each annotated mother centriole in the P7 serial scanning EM volume (right). Differentiatng cells in the immunofluorescent images were identified based on expression of P27^KIP^. Because the EM volumes lacks molecular markers we instead used cellular context and identified GCs near the EGL boundary as a pool of differentiating cells. Glia were included in the immunofluorescent analysis because SOX9 staining allowed us to distinguished them with confidence. (D) The length of each cilium from the widefield images is plotted on the left where each individual cilium measurement is represented by a small symbol and the average for each animal is represented by the larger symbol. The line and error bars represent the mean and standard deviation of the individual animal averages. The measured length of each cilium annotated in the P7 EM volume is quantified on the right.

GCs in the inner EGL have been reported to have fewer and shorter cilia than progenitors in the outer EGL (Ong et al., 2020). Cilia are also sparse in GC neurons as they mature (Chang et al., 2019). To examine cilia status on GCs throughout the entire process of neuronal differentiation and maturation, we analyzed cilia frequency and length throughout the cerebellar layers. We found that in the outer EGL ∼94% of the GC progenitors were ARL13B positive, indicating they had a cilium (Figure 1A, C). The fraction of ciliated cells decreased to 75% in the P27^KIP1^ positive differentiating cells in the inner EGL. Cells that had migrated beyond the EGL had dramatically fewer cilia: only 12% were ciliated in the ML and 10% in the IGL (Figure 1A, C). We also compared the length of cilia in each layer of the developing cerebellum (Figure 1D). The GC progenitor cilium averaged around 1 μm, while cilia in differentiating GCs were shorter. By contrast, glial cells (i.e., SOX9 positive cells) in the IGL layer had a mean cilium length of ∼3 μm (Figure 1C, D). Because we found that cilia were shorter in cells that had just begun differentiatiating and then became rare as GC neurons matured, we concluded that the short cilia on cells in the inner EGL disassemble, typically before GCs migrate into the ML.

While immunostaining enabled us to compare hundreds of cells from multiple specimens with molecular detail, many cilia measurements were close to the resolution limits of light microscopy, especially in z. We therefore employed volumetric EM to obtain more accurate measurements as well as detailed views of cilia ultrastructure. We uploaded, annotated, and analyzed centrosomes and cilia in GCs of the published 1.7 x 10^6^ mm^3^ (2,513 z slices) serial scanning EM data of a P7 mouse cerebellum (Wilson et al., 2019), which has 4 nm resolution in x and y, and 30 nm resolution in z. Just as with the immunofluorescent analysis above, we inferred a pseudo-timeline of GC differentiation by comparing cells across the cerebellar layers (Wilson et al., 2019). Without a molecular marker like P27^KIP1^, we could not discern the boundary between proliferating GC progenitors and GCs newly committed to differentiation (i.e., distinguish outer from inner EGL). However, we knew that all GCs that had reached the inner edge stained with P27^KIP1^ (Figure 1A), so we annotated the subclass of GCs close to the ML as “EGL boundary” (Figure 1B). A second difference between the two datasets was that the improved resolution in the EM data made it possible to identify the migrating GCs in the PCL (Figure 1B).

To evaluate cilia length and frequency modulation with the improved accuracy possible from EM, we traced and annotated cilia throughout the layers of the developing cerebellum. Detailed measurements for each cilium are included in Supplemental Table 1. We quantified the GC cilia frequency and length measured in each layer of the EM volume and compared the results to measurements from light microscopy. Overall, the cilia distributions were remarkably similar (Figure 1C, D). It is likely that the measured frequency of cilia in the EGL differs between the two measurements because the EGL in the EM data includes cells in both the inner and outer EGL. Cilia frequency was decreased in cells that had migrated to the ML, the PCL, and the IGL. When compared to the rest of the EGL, the GCs in the EGL boundary had similar cilia frequency, however, the median cilium length was much shorter (0.3 μm in the EGL boundary compared to 1.5 μm in the EGL). The few cilia we did observe in the IGL were typically short. However, two cilia in the IGL had lengths greater than 3 μm (Figure 1D). Many arriving immature GCs can lack synapses in the IGL, which can make them difficult to discriminate from glia/astrocytes in EM images. Therefore, we suspect these cilia belong to velate astrocytes (Farmer et al., 2016; Sun et al., 2017; Vong et al., 2015), which our immunofluorescence studies using glia molecular markers showed had significantly longer cilia than GCs in the IGL (Figure 1 D). Based on the reductions in cilia frequency, we conclude that cilia are largely absent from mature GC neurons in the IGL.

To assess the heterogeneity of cilia length we graphed the distribution of cilia lengths from all GCs in the EM dataset (Sup Figure 1A). We observed a bimodal distribution indicating two populations of cilia. Short cilia peaked around 300 nm and were all less than 750 nm. The distribution of longer cilia ranged from 1-2 μm. Long cilia were predominantly found in progenitors and cilia were shorter in newly differentiating GCs.

### Cilia on differentiating GCs are assembled post-mitotically and then disassemble

To further investigate timing and location of cilia disassembly during differentiation, we evaluated cilia length and frequency in a subpopulation of GCs that had just begun differentiating. To do this we injected P5 or P7 pups with BrdU and harvested the developing cerebella after either 12 or 48 h. BrdU is a thymidine analogue incorporated into DNA during S phase. Based on cell cycle dynamics in GC progenitors (Espinosa and Luo, 2008; Ho et al., 2020), differentiating GCs immediately exiting the cell cycle would be captured by the 12 h pulse-chasing. The 48 h chase permitted another round of division for cycling GC progenitors but also captured post-mitotic GCs that began differentiating and migrating immediately after the BrdU pulse (Figure 2A). We imaged the sections after staining for BrdU, P27^KIP1^ and ARL13B. All the BrdU labelled cells were in the EGL after 12 h chase (Figure 2B). About 30% of BrdU labelled cells also expressed P27^KIP1^, indicating that they exited the cell cyle after completing mitosis and had migrated to the inner EGL (Figure 2C). As expected, the number of BrdU positive cells in deeper layers increased with the 48 h chase (Figure 2B,C).

**Figure 2.**
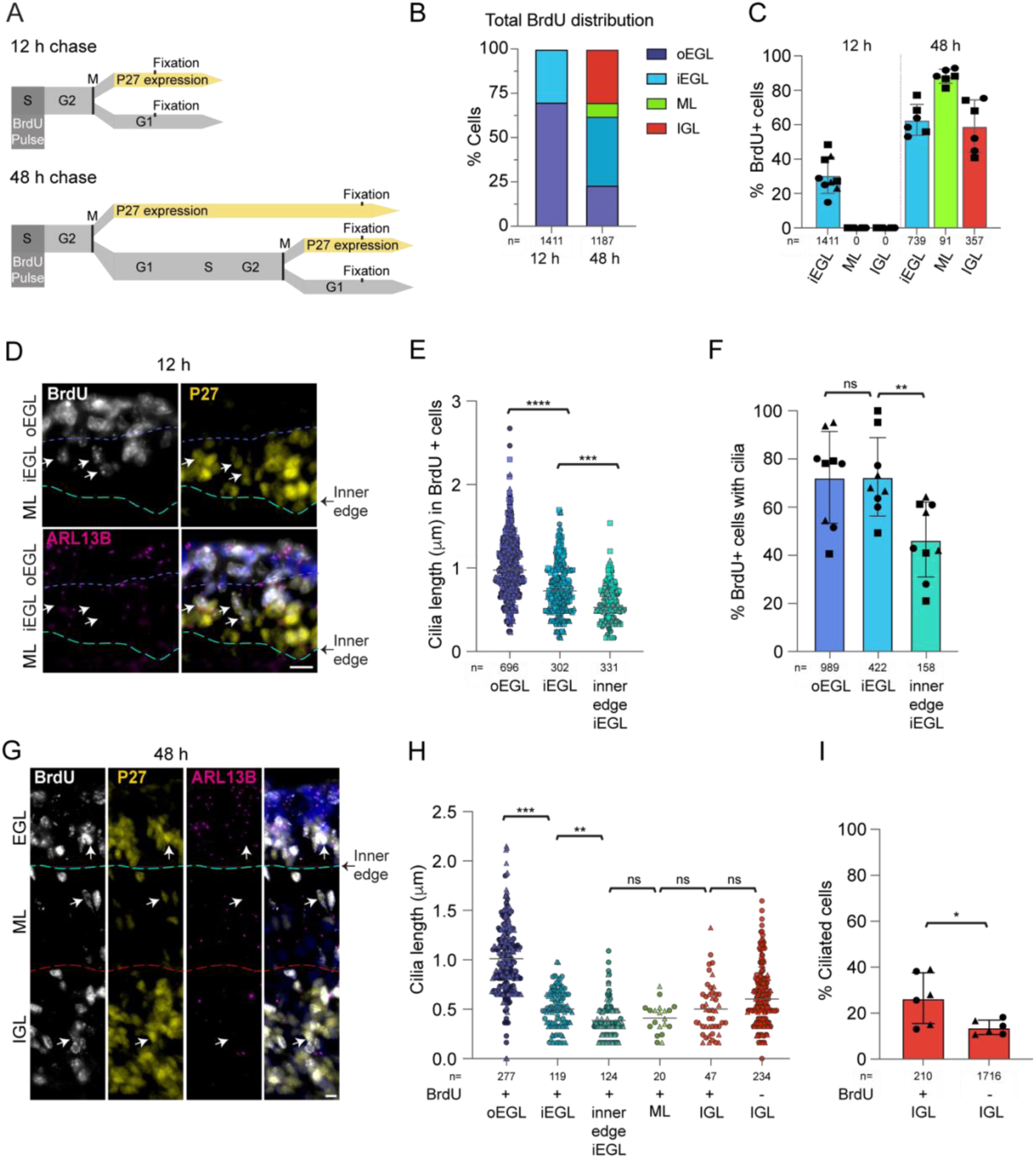
BrdU labeling reveals that early differentiating GCs have short primary cilia that subsequently disassemble. (A) P7 mice were injected with the thymidine analogue BrdU and cerebella were harvested after 12 h or 48 h. BrdU incorporates into DNA of cycling cells during the S phase. BrdU labeled cells that began differentiating became P27^KIP^ positive and migrated deeper. (B) BrdU labeled cells were classified by location in the developing cerebellum. The distribution of BrdU labeled GCs was quantified 12 h and 48 h after BrdU injection. (C) The percentages of total BrdU labeled cells in each layer that expressed P27^KIP^ 12 and 48 h after BrdU injection were quantified from 3 sections from 2-3 animals. Shapes denote biological replicates. (D) Sagittal sections of P7 mouse cerebella harvested 12 h after BrdU injection were immunostained with antibodies to BrdU (white), P27^KIP^ (yellow), ARL13B (magenta) and counterstained with Hoechst (blue in merge panel) before imaging with widefield microscopy. The approximate boundaries between layers are indicated in white and yellow. Arrows denote BrdU labeled P27^KIP^ positive cells. Scale bar: 10 μm. (E) BrdU labeled ciliated cells in the inner and outer EGL were distinguished based on P27^KIP^ expression and cells at the inner edge of the EGL were directly adjacent to the ML. The measured lengths of cilia in these cells is plotted. n= number of cilia counted. (F) The percentages of BrdU positive GCs with cilia in the outer EGL (P27^KIP^ negative), the inner EGL (P27^KIP^ positive), and at the inner edge were quantified 12 h after BrdU administration. (G) Sagittal sections of P7 mice cerebella harvested 48 h after BrdU injection were immunostained with antibodies to BrdU (white), P27^KIP^ (yellow), ARL13B (magenta) and counter-stained with Hoechst (blue in merge panel) before imaging using widefield microscopy. The approximate boundaries between layers are indicated in yellow. Arrows denote BrdU labeled P27^KIP^ positive cells. Scale bar: 10 μm. (H) The length of each cilium in the indicated layers 48 h after BrdU labeling is plotted. (I) The percentage of ciliated GCs in the IGL is plotted for BrdU labeled and unlabeled cells 48 h after injection. P values: 0.0.332 (*), 0.0021 (**), 00002 (***), <0.0001 (****).

To determine whether cilia shorten and/or disassemble as GCs progress through differentiation, we compared the cilium length and frequency at different stages of differentiation based on the migration through the cerebellar layers. GC cilia in the P27^KIP1^-expressing, BrdU positive cells of the inner EGL were shorter than the BrdU labeled progenitor cells in the outer EGL (0.7 μm and 1 μm respectively; Figure 2D-E). The BrdU and P27^KIP1^ positive cells directly at the inner edge of the EGL, poised to migrate across the ML, were shorter (0.5 μm) and less frequent than the P27^KIP1^-expressing, BrdU positive cells of the inner EGL that had not migrated as far (Figure 2E,F). Similar differences in cilia length and frequency were measured 48 h after BrdU treatment (Figure 2G,H). These results indicate that the small cilia shorten before arriving at the EGL boundary.

Although most GCs have no detectable cilium after migrating beyond the EGL, a small number of GCs in the IGL were ciliated. To determine whether cilia disassembly can continue outside the EGL we used the BrdU labeling to distinguish between newly arrived GCs and more mature neurons. Specifically, cells that express P27^KIP1^ but do not stain with BrdU arrived in the IGL before the BrdU positive GCs. Cilia lengths in the two populations were comparable (Figure 2H). However, we found that the cilia frequency was higher for BrdU positive cells in the IGL (Figure 2I). Together, these data indicate that cilia in differentiating GCs shorten in the inner EGL and that cilia continue to be disassembled in maturing GCs in the IGL.

### Cilia disassembly in progenitor GCs is distinct from disassembly in differentiating GCs

Cilia on post-mitotic GCs at the EGL boundary were less than 750 nm (Figure 1D). This population of short cilia are unlikely to arise from gradual shortening of longer cilia in the outer EGL because cilia length distribution in GCs was not continuous but was bimodal, peaking around ∼0.5 and ∼1.5 μm, with very few intermediate length cilia (Sup. Figure 1A). Light microscopy has shown that cultured mouse medulloblastoma GCs resorb cilia prior to mitosis *in vitro* (Ho et al., 2020). There are thus two ways cilia disassembly during progenitor cell division could have generated the population of short cilia: 1) each cilium was resorbed prior to GC progenitor mitosis and cilium regrowth was stunted in differentiating daughter cells; or 2) short cilia were remnants of progenitor cilia retained though mitosis as has been observed in other neural progenitors (Paridaen et al., 2013). To determine if any remnants of cilia remain associated with mother centrioles during mitosis *in situ*, we examined centrioles in dividing progenitor cells in the P7 EM volume. We first identified progenitor cells in S-phase, G2, and mitosis. Procentrioles were visible in the EGL templating off the mother and daughter centrioles indicating the cells were in S phase and G2 (Figure 3C) (Kumar and Reiter, 2021; O’Toole and Dutcher, 2014). Although some procentrioles could be classified as nascent or mature, the distinction was sometimes complicated by cut angle and imaging artifacts, so for analysis we grouped all cells with a procentriole into a combined S/G2 category. We found cilia in most S/G2 progenitor cells (Figure 3A-C) indicating that cilia disassembly occurs just before cells enter mitosis. This is similar to results in cultured GC progenitors and other cells, where cilia disassemble sometime after reaching S phase (Ford et al., 2018; Ho et al., 2020). Generally, cilia length in S/G2 cells was similar to cells in G1/G0 (Figure 3B,C). Cilia disassembly appeared to involve internal intermediates including late-stage centriole-associated membranes (Figure 3C,D). Cilia disassembly was completed by the time GCs entered mitosis. We found no axonemes or membrane vesicles associated with mitotic centrioles (Figure 3A,E). The early ciliogenesis intermediates shown in Figure 3F were observed only in GCs in which cytokinesis had progressed sufficiently that midbody constriction was evident and the cytokinetic bridge was largely enveloped within a daughter cell. Based on these observations, we conclude that cilia are completely disassembled prior to mitosis and ciliary remnants are not carried through into differentiating GCs. Hence, the disassembly of short cilia during GC differentiation is distinct from pre-mitotic cilia disassembly. To distinguish the processes, we will refer to the novel dismantling of cilia in post-mitotic GCs as cilia deconstruction.

**Figure 3.**
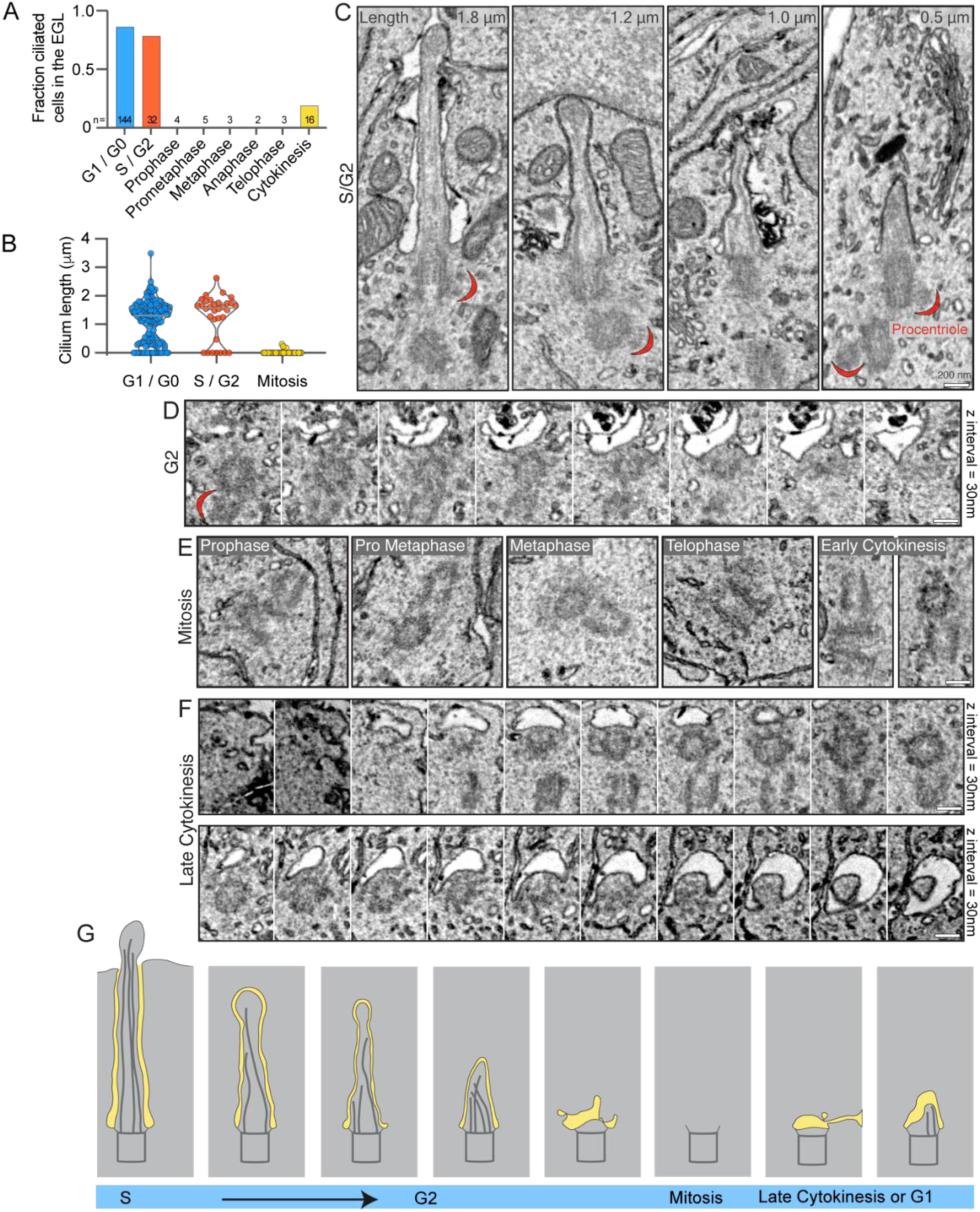
Cilia are completely resorbed during mitosis in GC progenitors. (A) The cell cycle status of each cell in the EGL of the P7 SEM volume was determined and annotated. The fraction of cells in each phase with primary cilia is quantified. (B) The length of each EGL GC cilium is plotted by cell cycle phase. G1 and G0 cells cannot be distinguished; both have two centrioles. (C) Serial scanning EM images of cilia from 4 individual cells with duplicating centrioles that indicate the cells are in S phase or G2. Cilium length is noted in the upper right corner and procentrioles visible in the same section are indicated with a red crescent. (D) Serial sections of a non-ciliated mother centriole and procentriole in a G2 cell in the EGL. (E) Representative SEM images of non-ciliated centrioles in GC progenitors in each stage of mitosis. (F) Serial sections of mother centrioles from late-stage cytokinesis cells with a ciliary vesicle (upper) or nascent cilium (lower). (G) Amodel of the process of cilia resorption using representations of the cilia presented in panels C-F. Prior to mitosis cilia disassemble, are absent during mitosis, and reappear late in cytokinesis.

### Most GC cilia are intracellular and can be concealed from the external environment

We next examined the ultrastructure of cilia within the volumetric EM to gain insights into differences between long and short cilia and in this process discovered that external exposure of cilia was also modulated. While cilia in some cells protrude directly from centrioles docked at the cell surface, most cilia that extend from centrioles deeper in the cell are ensheathed in a membranous pocket, called a ciliary pocket that invaginates from the cell surface and anchors at the distal appendages (Rohatgi and Snell, 2010). As has been seen in published images (Chang et al., 2019; Ong et al., 2020; Spassky et al., 2008), many progenitor GC cilia had ciliary pockets that we call “pocket cilia” (Figure 4A, Sup Video 1). However, we observed that only a small portion of the pocket cilia extended into the extracellular space; most of the cilia were hidden in the deep ciliary pocket (Figure 4A). The tip of over 25% of pocket cilia were, however, shrouded upon exiting the pocket because their tips were enveloped in a plasma membrane invagination of an adjacent cell (Supplemental Fig 2). We observed a second class of cilia that lacked a ciliary pocket that we call “surface cilia”. These cilia extended directly from a mother centriole docked at the plasma membrane and were, therefore, completey exposed (Figure 4B, Sup Video 2). The third and largest group of cilia were completely contained in an ensheathing membrane – submerged in the cytoplasm – without exposure to the extracellular environment (Figure 4C,D; Sup Videos 3,4). We refer to these as “concealed cilia” because their access to extracellular signals such as SHH was likely restricted. We conclude that, with the exception of surface cilia, membrane structures modulated cilia exposure to exracellular milieu.

**Figure 4.**
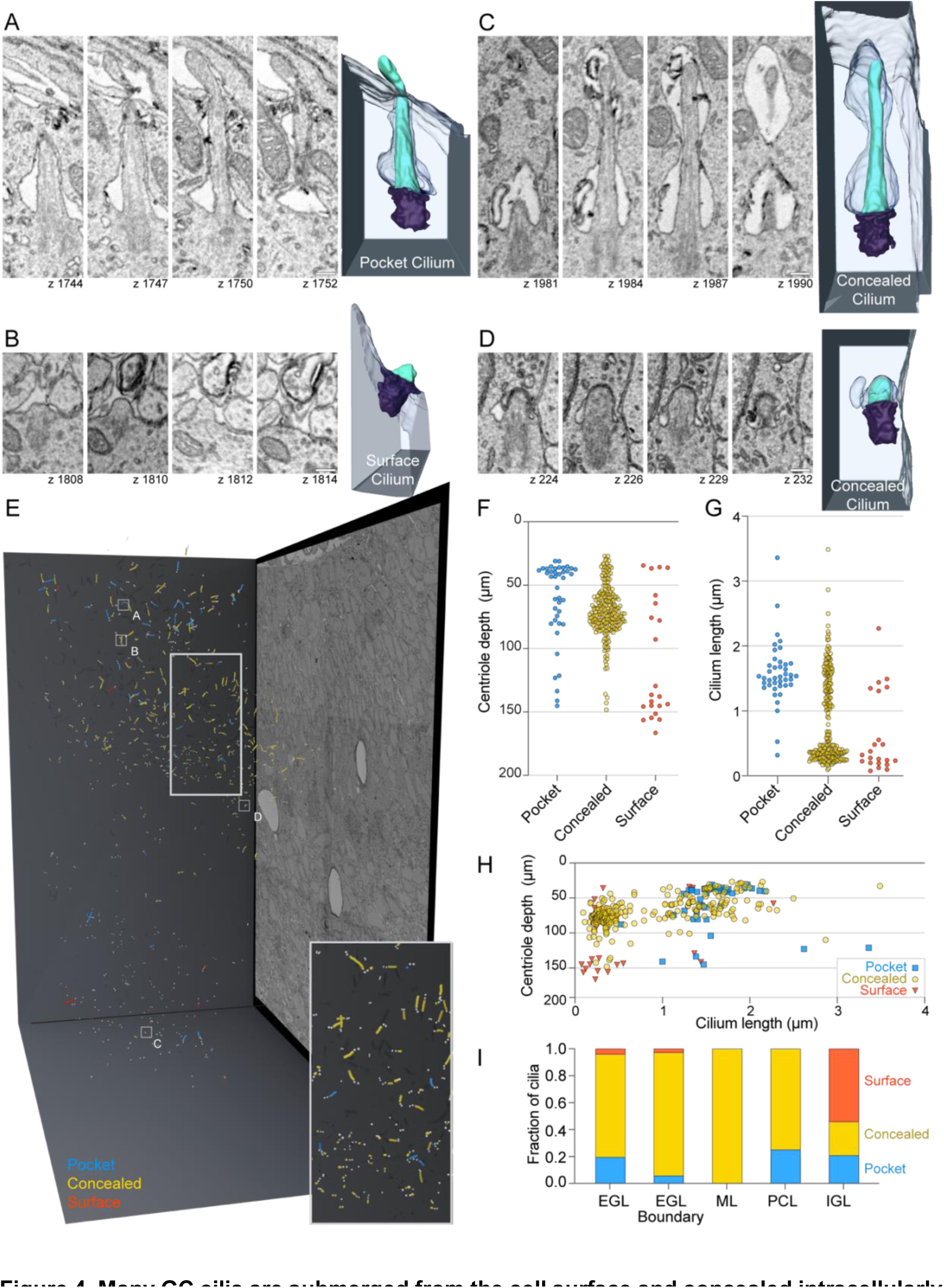
Many GC cilia are submerged from the cell surface and concealed intracellularly. (A – D) Representative pocket (A), surface (B) and concealed (C and D) cilia from P7 large serial scanning EM volume are displayed. Select z slices of each cilium are presented to the right of the 3D reconstructed image of each cilium. Scale bar: 200 nm. (E) Annotated cilia and centrioles are represented in the 3D space of the serial scanning EM volume. Annotated centrioles are represented as grey spheres and cilia as lines. Cilia color represents the classification of cilia as Pocket (blue), Concealed (yellowyellow), or Surface (red). The positions of the cilia displayed in panels A-D are indicated. (F) The depth of mother centrioles is plotted for each type of cilium. In (F) and (H) the y-axis is inverted such that higher depth values fall farther below the Y-axis. (G) Cilium length is plotted for each type of cilium. (H) The length of each annotated cilium is plotted as a function of mother centriole depth. The cilia are color coded as classified. (I) The fractional distribution of cilia in each layer is plotted and colored according to cilia type.

To determine whether any of the ciliary ultrastructures were enriched at specific developmental stages, we examined the distribution of pocket, surface and concealed cilia within the developing cerebellum. A 3D projection of all cilia by type is displayed in Figure 4E. The distributions of each cilium type by centriole depth (distance from pia), cilium length and layer are quantified in Figure 4 F-H and are shown with respect to the cell cycle in Sup. Figure 1 E-F. Concealed cilia were dominant in the EGL, both in G1/G0 and S/G2 cells. Pocket cilia were generally >1 μm and most were found in the outer EGL indicating that they were present on GC progenitors. Overwhelmingly, the cilia at the EGL boundary were both short and concealed. Surface cilia, while occasionally observed in the EGL, were the dominant cilium type in the IGL. Because cilia deconstruction occurred in both the EGL boundary and IGL, we conclude that the disassembling cilia during GC differentiation were most likely concealed and surface cilia.

### Large-scale EM reveals novel cilia disassembly intermediates associated with maturing GCs

To gain further insights into the process of cilia deconstruction in differentiating GC neurons, we closely examined cilium and centrosome ultrastructures within the EGL boundary. We recognized that a group of short, concealed cilia were distinct (Figure 5A and Sup Figure 3A compared to Figure 5B and Sup Figure 3B). Instead of tapering or rounding slightly at the tip like typical cilia, this subset had a rounded, bulbous shape. We labeled these as protein-rich cilia because the cilioplasm appeared more homogenous, amorphous, and darker than the adjacent cytoplasm or than the cilioplasm of other cilia, which indicated additional electron dense staining. Axoneme microtubules and transition zones (Reiter et al., 2012) were difficult to discern in this subset of cilia either because they had depolymerized or because they were indistinguishable in the darker staining cilioplasm. We plotted the depth of the protein-rich cilia in comparison to the more typical short, concealed cilia (Figure 5C). While neither was restricted to a single layer, both were largely found in the EGL boundary, suggesting that protein-rich cilia were deconstruction intermediates observed during GC differentiation, but not during pre-mitotic cilia resorption.

**Figure 5.**
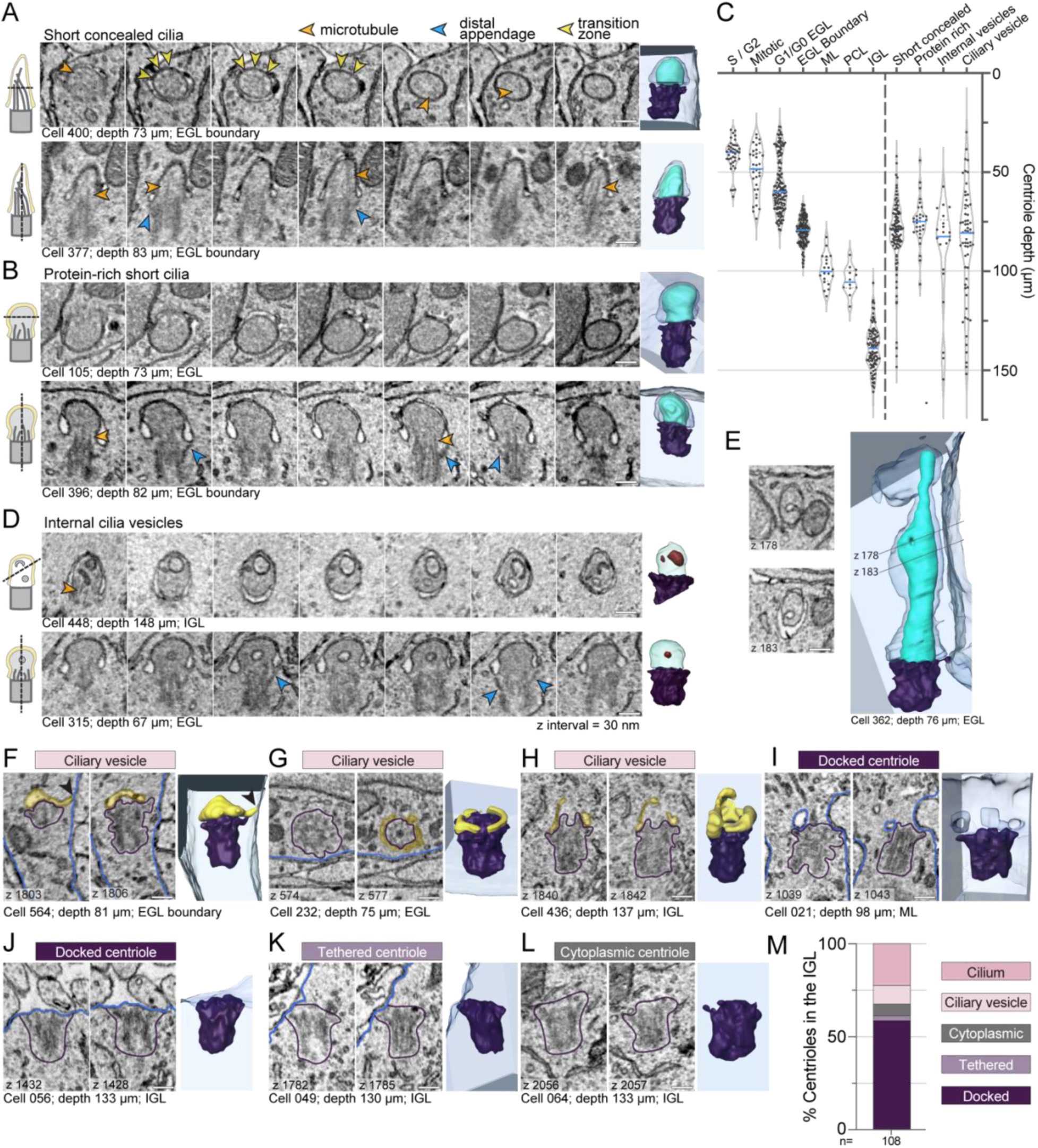
Large-scale EM reveals novel cilia disassembly intermediates in differentiating GCs. (A, B, and D) Serial sections of representative short, concealed cilia (A), protein-rich short cilia (B) and cilia with internal vesicles (D) are displayed. The section orientation is indicated by the cartoon on the left and the 3D reconstructions of each cilium is displayed on the far right. (C) Mother centriole depth is plotted on an inverted y-axis such that higher depth values fall farther below the Y-axis. To the left of the dotted vertical line, the centrioles are classified by layer, with the outer EGL the cells broken out by cell cycle phase. On the right side of the dashed line, the mother centrioles are grouped based on the type of cilia deconstruction intermediates observed. (E) Two representative EM sections showing invagination of the ciliary membrane are shown to the left of the 3D reconstruction of the concealed cilium from the EGL. (F – L) For each centriole, two representative EM sections are displayed to the left of the 3D reconstruction. The classification of the centriole is indicated as ciliary vesicle (F-H), docked centriole (I-J), tethered centriole (K), or cytoplasmic centriole (L). On the EM images, yellow transparent overlay indicates ciliary vesicles, blue highlights the cell boundary and purple encircles the centriole. (K) The percentage of GCs in the IGL with centrioles in each classification are quantified. All scale bars are 200 nm.

We searched for evidence of vesiculation from cilia or ciliary severing, established processes that shorten cilia (Cao et al., 2015; Esparza et al., 2013; Mirvis et al., 2019; Nager et al., 2017; Phua et al., 2017; Wang and Barr, 2016; Wang et al., 2015; Wood and Rosenbaum, 2014; Wood and Rosenbaum, 2015). We found only one possible example (Sup Fig 3 C,D). It was a concealed cilium with a constriction just above the base. If severing occurred the cilium would not be shed, but rather enclosed in a large intracellular vesicle. The infrequency of severing related structures indicates that more often short cilia arise from limited cilium growth in differentiating cells, and that disassembly in differentiating GCs involves a cilia deconstruction process, not cilia severing.

Upon examining cilia ultrastructure we also noted that a subset of cilia contained internal vesicles (Figure 5D). These cilia were largely found in the EGL boundary; however, some cilia with internal vesicles were present in the more mature GCs of the IGL. Occassionally protein-rich cilia contained internal vesicles. It seemed plausible that internal vesicles could have entered the cilium from the base, however, we found cilia with invagination of the ciliary membrane (Figure 5E), suggesting that internal vesicles were derived from invagination of ciliary membrane. Endocytosis of ciliary membrane could be a disassembly strategy that reduces the surface area of the cilium. While internal vesicles have been previously reported in chondrocytes (Jensen et al., 2004) and in specialized primary cilia, such as in olfactory neurons and in photoreceptors (Chuang et al., 2015; Jana et al., 2018; Reese, 1965), we are not aware of prior association with cilium disassembly.

The volumetric EM also provided unique views of centrioles in differentiating GC neurons. Throughout the cerebellar layers, but especially enriched in EGL boundary, we found membrane structures associated with the mother centriole distal appendages (Figure 5A, F-I). Figure 5F (Sup Video 5) shows a basal body in a GC at the EGL boundary capped with a ciliary vesicle and a short tube extending to the cell surface. During ciliogenesis, similar dynamic tubules project to the cell surface (Ganga et al., 2021; Insinna et al., 2019; Stuck et al., 2021). The structures in Figure 5G and H resembled pre-ciliary toroid membranes (Sup Videos 6,7) (Ganga et al., 2021; Insinna et al., 2019; Stuck et al., 2021); however, the membrane in Figure 5H extended up in places like a ciliary membrane. Based on depth, we inferred which centrioles were likely in cycling progenitor cells and which were disassembly intermediates in differentiating cells (Figure 5C). We also found unciliated centrioles that had invaginations of plasma membrane anchored to distal appendages (Figure 5I, Sup Video 8, and Sup Figure 3). We conclude that distal appendage-associated membranes in differentiating GCs in the EGL boundary, ML, PCL, and IGL are late-stage intermediates during cilia disassembly.

As we examined the centrioles in the IGL we made another unanticipated discovery. We found unciliated mother centrioles anchored directly to the plasma membrane by their distal appendages. Most of these centrioles were fully docked, similar to the basal body of a surface cilium (Figure 5J, Sup Video 9), and a few were tethered by association of just a few distal appendages (Figure 5K, Sup Video 10). Only a small subset of centrioles were immersed in the cytoplasm without any ciliary vesicles (Figure 5L, Sup Video 11). The distribution of ultrastructures found in the IGL is graphed in Figure 5M. The high frequency of docked centrioles suggests docked centrioles are a hallmark end-state of cilia deconstruction during GC neurogenesis.

### Unciliated mother centrioles dock at the plasma membrane in adult mouse GC neurons

A prediction of the pathway for cilia deconstruction described above is that it results in absence of cilia in mature GC neurons. We therefore examined the ciliation state by immunoassaying sagittal sections of adult (P25) mouse cerebella with antibodies to ARL13B. In the adult, the EGL no longer exists and GC neurons pack densely in the IGL. Purkinje neurons and glia are also present in both the PCL and the IGL and can be ciliated. Therefore, like in the developing cerebellum, we used antibodies to P27^KIP1^ and SOX9 to positively identify GCs and glia, respectively (Farmer et al., 2016; Sun et al., 2017; Vong et al., 2015) (Figure 6A). We found that in the IGL <1% of GC neurons were ciliated, whereas 70% of glia (ie. SOX9+ cells) were ciliated (Figure 6A, B). The cilia in the glial cells averaged ∼4 μm. By contrast, GC cilia averaged less than a micron in length by widefield fluorescence microscopy (Figure 6C). To determine whether the unciliated centrioles in adult GC neurons are docked, we utilized two recently generated EM volumes created to investigate synaptic connections (Nguyen et al., 2023). We located, annotated and analyzed more than 100 centrosomes in each of two datasets of serial-section transmission EM from lobule V and lobule X of the adult mouse vermis imaged at a resolution of 4.3 × 4.3 nm^2^ per pixel ((Nguyen et al., 2023); unpublished). Almost every mother centriole was attached to the plasma membrane through adhesion of the distal appendages (Figure 6 E-G). We only found a single, short cilium with internal vesicles in the lobule V volume (Figure 6F, Supp. Figure 4A) and 5 tethered and 3 cytoplasmic centrioles in the lobule X volume (Figure 6F). In combination with the analysis of docked centrioles in the P7 IGL, we conclude that GC neurons in adult mice have largely lost their cilia by a pathway involving cilia deconstruction and centriole docking during GC neural differentiation.

**Figure 6.**
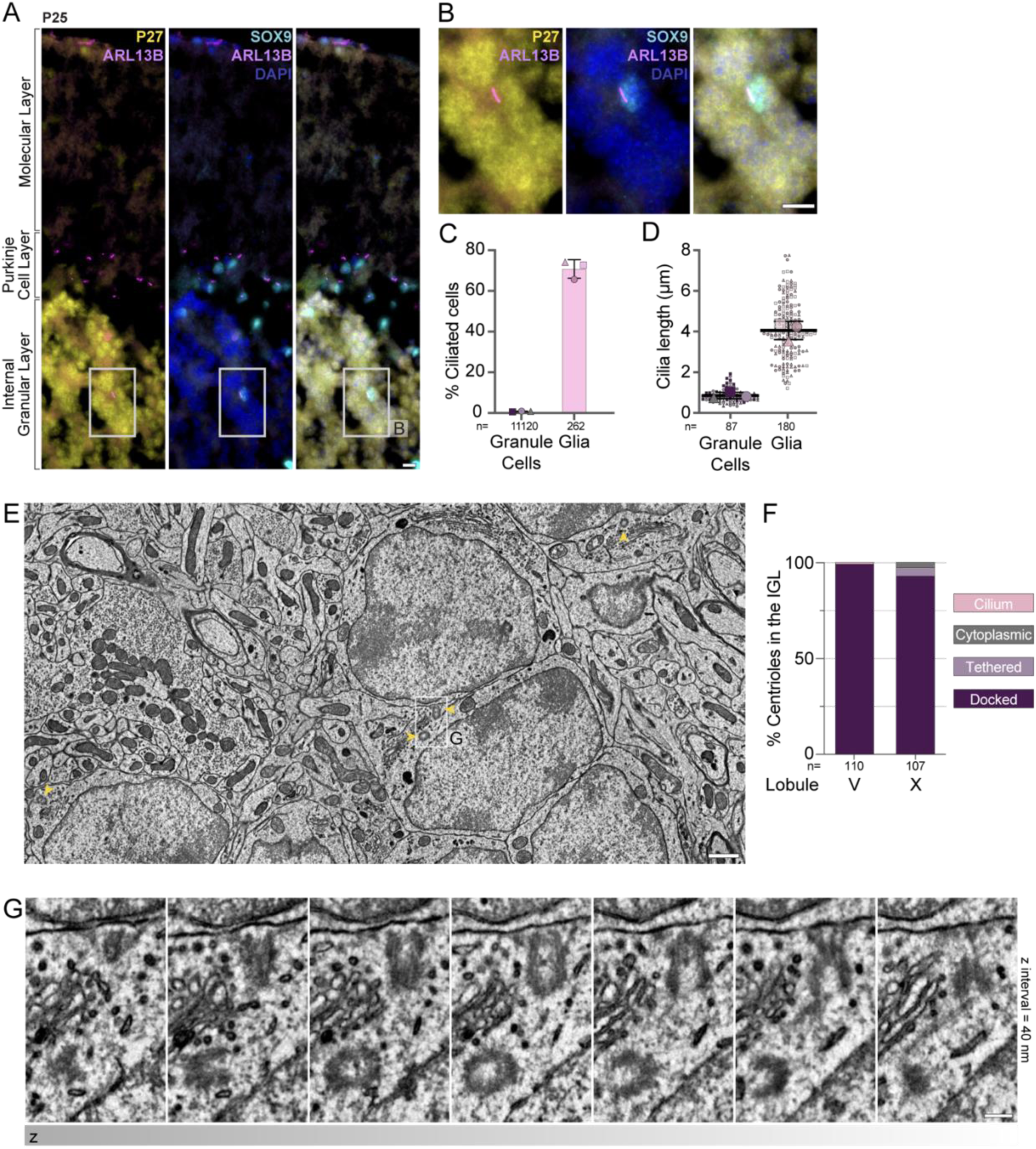
Cilia are absent in mature GC neurons despite docking of mother centrioles at cell surface. **(A)** Sagittal cerebella sections from a P25 mouse were immunostained with antibodies to ARL13B (magenta) to visualize cilia, P27^KIP1^ (yellow) to mark GC neurons, SOX9 (cyan) to mark glial cells, and counterstained with DAPI (blue) before imaging with widefield microscopy. Cilia are prominent in SOX9 glial cells; however, they are not detected on P27^KIP^ positive GC neurons. Scale bar: 10 μm. **(B)** Enlarged view of granule cells and a glial cell from the IGL of panel A. **(C** – **D)** Cilia frequency (C) and length (D) were measured in immunostained images acquired with widefield microscopy. Average frequencies of 3 sections each, from 3 animals is shown in C and individual cilia measurements from the same sections are shown as small symbols in D with the average per animal represented by the large symbol. The line and error bars represent the mean and standard deviation of the individual animal averages. **(E)** EM image from a serial-section transmission EM volume of the IGL of an adult mouse cerebellum. Centrioles are indicated with yellow arrowheads. Scale bar: 1 μm. **(F)** The percentage of GCs in the adult IGL with centrioles in each classification are quantified. **(G)** Serial sections of the centriole highlighted in (E). Scale bar: 200 nm.

We investigated the cellular context of the docked centrioles in search of functional insights. Centrioles docked at the plasma membrane in the small region of each GC in areas where Golgi, mitochondria, and most other organelles were concentrated. Ciliary rootlet structures were visible in many GCs. We measured the orientation of centrioles and found no positioning or orientation bias within the tissue (Supp Figure 4C). We also determined the type of structures extracellular to each mother centriole. Unlike T cells which transiently dock mother centrioles at the immune synapse to direct release of lytic granules toward a target cell (Stinchcombe et al., 2015), we found diverse structures opposite the GC docked centrioles including the soma of other GCs, axons, dendrites, and glial processes including myelin sheaths (Supp Figure 4C) and no evidence of centriole nucleated microtubules forming highways positioned to deploy vesicles. Instead of being oriented relative to external factors, centriole docking appeared to influence internal organization of the GC soma. Although centriole docking can be the first stage in the biogenesis of surface cilia, during differentiation, GCs disassemble cilia, anchor the centriole to the plasma membrane, and remain unciliated.

## DISCUSSION

We report here an almost complete absence of primary cilia in adult GC neurons in the cerebellum. The lack of cilia in GC neurons is remarkable because most other neuronal subtypes have cilia (Green and Mykytyn, 2014; Louvi and Grove, 2011; Ott et al., 2023; Wu et al., 2023). It is known that cilia are required for proliferation of GC neural progenitors (Chizhikov et al., 2007; Spassky et al., 2008). We discovered that primary cilia were present early in differentiation, disassembled as GC neurons matured and did not regrow despite mother centrioles docking like basal bodies at the cell surface. We detected previously undescribed disassembly intermediates that parallel structures observed during ciliogenesis (Ganga et al., 2021; Insinna et al., 2019; Stuck et al., 2021), To our knowledge, this is the first description of cilia disassembly in post-mitotic, differentiating cells. To distinguish it from pre-mitotic cilia resorption, we refer to cilia disassembly in differentiating GCs as cilia deconstruction.

The concealed cilia we observe could be intermediates in the internal ciliogenesis pathway (Sorokin, 1962; Sorokin, 1968) or could be reversibly generated from surface exposed cilia, as reported in cultured cells (Rivera-Molina et al., 2021). The prevalence of concealed primary cilia could not have been predicted from light microscopy images. Also remniscent of intracellular ciliogenesis intermediates, we found that some centrioles in maturing GC neurons had distal appendage associated membranes (Breslow and Holland, 2019; Zhao et al., 2022). However, we were able to infer that the observed cilia intermediates were disassembling, not assembling, due to their location and the correlation between cell position and developmental status in the cerebellum. The presence of concealed cilia and the observed similarities suggest that the mechanisms used to build and maintain cilia might influence deconstruction of pre-formed cilia.

The shapes of centriole docked membranes were especially diverse, possibly due to membrane dynamics not captured in the single EM volume. Unlike in pre-mitotic cells, we were unable to assemble a single linear disassembly pathway from the observed disassembling structures in differentiating GCs. We propose that the heterogeneity of the observed intermediates reflected diversity in the cilia deconstruction and centriole docking pathway. In Sup Figure 5 we have illustrated four possible progressions of cilia deconstruction which include different observed intermediate structures that could all resolve as docked centrioles. Several hypothesize that membrane pores or tubules, similar to the dynamic PACSIN and EHD1 dependent tubules involved in ciliogenesis (Ganga et al., 2021; Insinna et al., 2019; Stuck et al., 2021), join the internal cilia membrane to the plasma membrane. The difference between each model is the extent of cilia disassembly prior to centriole docking. In the first, fusion of the ciliary sheath before complete cilia deconstruction results in surface cilia similar to those observed in the IGL (many of which had atypical morphology, lacked axonemes, or contained intraciliary vesicles). The second and third illustrations involve fusion of distal appendage anchored vesicles to the plasma membrane resulting in plasma membrane invaginations that would be resolved to generate docked centrioles. These possibilities can be compared to the fourth proposed progression: the conventional paradigm of a cytoplasmic mother centriole docking directly to the plasma membrane. Live monitoring of cilia deconstruction and centriole docking will be necessary to determine if one or all of these docking pathways occur within the tissue.

The newly described cilia deconstruction pathway appears to include different intermediates than those seen in ciliary disassembly that occurs prior to cell division (Pugacheva et al., 2007; Tucker et al., 1979) or during the post-fertilization stage of *Chlamydomonas* (Pan and Snell, 2003; Pan and Snell, 2005; Pan et al., 2004). Thus, our results suggest that established cilia disassembly mechanisms (Liang et al., 2016; Malicki and Johnson, 2017; Wang and Dynlacht, 2018) may not fully account for cilia loss during GC differentiation. We find no evidence for rapid disassembly of microtubules (Pan and Snell, 2005; Pugacheva et al., 2007), internalization of a portion of the axoneme (Rieder et al., 1979), or shear stress (Liang et al., 2016), which have all been shown to contribute to cilia disassembly in other systems. The persistence of a few cilia in postnatal IGL GCs beyond the 48 h BrdU pulse indicates a wide spatiotemporal distribution of deconstructing cilia during GC differentiation. The process of cilia deconstruction involves gradual axoneme depolymerization and recovery of the ciliary membrane, as suggested by enhanced electron-dense staining in protein-rich cilia and internal cilia vesicles. In addition, the observed late stage centriole-associated vesicles indicate that ciliary membrane and ciliary pocket retrival are coordinated in the final stages. Cilia loss during differentiation is not unique to GC neurogenesis. Adipocytes (Hilgendorf et al., 2019; Zhu et al., 2009), retina pigment epithelial cells (Patnaik et al., 2019), myoblasts (Fu et al., 2014), steroidogenic adrenal cortical cells (King et al., 2009; Mateska et al., 2020) and oligodendrocytes (Buchanan et al., 2022) are all derived from ciliated progenitor cells. Hypertrophic chondrocytes also have lower ciliation compared to the columnar chondrocyte precursors (Hwang et al., 2018). Cilia absence could be caused by prevention of cilia re-growth after pre-mitotic cilia resorption; however, in some of these diverse contexts, the cilia deconstruction pathway we identified may be responsible for cilia loss.

Developmental and circadian regulated cilia disassembly followed by regrowth has been reported in diverse systems (Das and Storey, 2014; Lepanto et al., 2016; Toro-Tapia and Das, 2020). During GC neurogenesis, however, not only did the mother centrioles in adult granule cells remain unciliated, they docked at the plasma membrane. Such docking was unexpected because there is only few systems where centriole docking without cilia extension has been described that includes the immune synapse (Stinchcombe et al., 2015). Mother centrioles in T cells transiently dock at the plasma membrane adjacent to the target cell. Unlike docked centrioles at the T cell synapse, docked cerebellar GC centrioles do not appear to create a hotspot for exocytosis. Because centriole docking is pervasive in adult GC neurons, it is likely that centriole docking is permanent. We found additional evidence for centriole docking in historic electron micrographs of GC neurons (Del Cerro and Snider, 1969) and gamma cells in the adrenal cortex (Wheatley, 1967).

The pathway of cilia concealment and deconstruction described here may be important to prevent aberrant SHH signaling. Soluble SHH secreted by Purkinje neurons permeates the EGL, yet upon onset of differentiation, GCs stop responding to the mitogenic signal (Corrales et al., 2004). Concealment of cilia in the cytoplasm could ensure that proliferative programming is not reactivated during development by SHH detection but active GLI-mediated repression by cilia is maintaned (Han et al., 2009; Kopinke et al., 2020). Deconstructing cilia may also be important in mature tissue. One prevalent subtype of medulloblastoma, a cerebellar brain tumor, is caused by unrestricted proliferation of GCs with progenitor characteristics (Schuller et al., 2008; Yang et al., 2008) from aberrant activation of the SHH signaling pathway (Oliver et al., 2005; Pak and Segal, 2016; Shimada et al., 2018). The GCs in SHH-medulloblastoma can be ciliated (Di Pietro et al., 2017; Han et al., 2009; Youn et al., 2022). In mice embryos, a docked centriole is sufficient to provide cues to maintain renal tubule architecture but postnatally a cilium is required for tubular homeostasis (San Agustin et al., 2016). However, lack of cilia formation from the docked centrioles in adult GCs likely suppress SHH receptivity, blocking proliferative potential and dedifferentiation. Our results suggest that both proper deconstruction of cilia and prevention of cilia regrowth is needed to permanently disable SHH signaling.

In the accompanying manuscript, we investigated the molecular changes that accompanied GC differentiation in accomplishing cilia deconstruction and what prevents cilium growth from docked mother centrioles in mature GCs (Constable, Ott et al. co-submitted). We find that global, developmentally programmed, diminution of cilium maintenance, rather than active disassembly processes, caused cilia deconstruction in differentiating GCs. Furthermore, binding of centriolar cap proteins to the docked mother centrioles prevented ciliation in adult GC neurons.

## Acknowledgements

This project was funded by the National Institutes of Health (1R35GM144136 to S.M,, R21NS085320 and RF1MH114047 to W.C.A.L., and 1S10OD028630 to Microscopy Core Facility in UT Southwestern), an Alex Lemonade stand Foundation A-Award to S.M., and the Howard Hughes Medical Institute. The authors would like to acknowledge the Quantitative Light Microscopy Core, a Shared Resource of the Harold C. Simmons Cancer Center in UT Southwestern, supported in part by an NCI Cancer Center Support Grant, 1P30 CA142543-01. The content is solely the responsibility of the authors and does not necessarily represent the official views of the National Institutes of Health. We thank molecular pathology and mouse animal care facility in UT Southwestern. We also thank Tom Kazimiers from kazmos GmbH for CATMAID support and Will Patton and David Ackerman from Janelia Scientific Computing for supporting code to extract quantitation. Henrique Ludwig helped annotate centriole vectors. We also thank Andy Moore for discussions about figures and Christina Gladkova, Lauren Porter, Mol Mir, Lorena Bendetti, Chris Obara and Cayla Jewett. for helpful discussions and comments on the manuscript. This collaboration originated as a result of discussions at a BSCB GenSoc UK Cilia Network e-symposium.

## Author contributions

C.M.O., S.C. and S.M. conceived the project, designed experiments, and analyzed most of the data. C.M.O., S.C., J. L.S. and S.M. wrote the paper with inputs from all authors. S.C. and K.W performed immunofluorescence experiments. C.M.O annotated and analyzed EM data. T.M.N and W.A.L. provided EM datasets of adult mice.

## Materials and Methods

### Mouse handling and genotyping

All animal studies were approved in accordance with UTSW Institutional Animal Care and Use Committee regulations and were conducted in accordance with NIH guidelines for the care and use of laboratory animals. CD1 mice were purchased from Charles River laboratories and maintained under standard conditions. Mice were treated with 10mg/ml BrdU (Bromodeoxyuridine/ 5-bromo-2’-deoxyuridine) dissolved in PBS using intraperitoneal injection at 50mg/kg.

### Mouse brain processing

Mice were procured at the appropriate age and fixed by trans-cardial perfusion using 4% paraformaldehyde (PFA) in PBS after appropriate anesthesia for their age (either isoflurane or cold exposure on ice) according to IACUC regulations. Brains were removed and further fixed in 4% PFA/PBS overnight at 4oC on a rotator, then immersed in 30% sucrose in PBS until brain sank to the bottom of the tube (∼48 h). Brains were cut in half in the sagittal direction and embedded cut face down in cryomolds using OCT embedding media (BioTek, USA) and frozen on dry ice until solid. Blocks were stored at -80oC until sectioning on a Leica Cryostat model CM1950 cryostat at 15-30 μm thickness. Sections were stored at -20°C or -80°C until staining.

### Immunofluorescence staining and light microscopy

Sections were thawed at room temperature and OCT was removed by immersion in PBS. Sections were blocked using 3% serum (donkey) in PBS with 0.3% Triton-X 100 for 30mins. Primary antibodies were diluted in blocking solution at the appropriate dilutions and incubated overnight at room temperature in humid chamber. Primary antibodies: ARL13B (1:1000, UC Davis/NeuroMab #75-287), BrdU (1:500, Abcam #ab6326), P27^KIP^ (1:400, BD Biosciences) #610241), and SOX9 (1:500, Millipore #ABE571). For BrdU staining we performed 2N HCl pre-treatment for 15 min at 37°C before blocking solution was applied. Sections were incubated with the indicated isotype specific secondary antibodies for 2 h at room temperature. Sections were washed three times, DAPI (1μg/ml) Sigma) was included in the final wash to stain nuclei. Stained tissues were mounted using Fluromount-G (Southern Biotech) and allowed to dry overnight. Stained slides were imaged within 2-3 days and stored at 4°C (short term) or -20°C (long term).

Images were acquired on a widefield microscope (AxioImager.Z1; ZEISS), confocal microscope (Zeiss LSM880) or a spinning disk confocal microscope (Nikon CSU-W1 SoRa). Images in the widefield microscope were acquired using a Plan Apochromat objective (40×/1.3 NA oil and 63×/1.4 NA oil) and sCMOS camera (PCO Edge; BioVision Technologies) controlled using Micro-Manager software (University of California, San Francisco) at room temperature. Images in the confocal microscope (Zeiss LSM880) were acquired using Plan Apochromat objective (63×/1.4 NA oil). Images in the spinning disk confocal microscope (Nikon CSU-W1 SoRa) were acquired using a Plan Apochromat objective (100×/1.45 NA oil), a sCMOS camera (Hamamatsu Orca-Fusion), and a Piezo z-drive for fast z-stack acquisition controlled using Nikon NIS-Elements software at room temperature. Between 10 and 30 z sections at 0.2 µm intervals were acquired.

### Image analysis

#### Cilia length and number determination

Images were analyzed using FIJI (Schindelin et al., 2012). Different cerebellum layers were identified using nuclei, and P27 markers. Cilia length was manually determined by tracing in each zone at zoom level 200-300% using the freehand draw tool, and measurement recorded using measure tool. Cilia already traced were permanently marked with draw tool to ensure unique cilia were measured when moving around the image. Zones were completed in their entirety before moving on to the next zone. The number of cilia was determined by counting the number of cilia measured. The number of cells was determined by counting the total number of nuclei found in each section as stained by Hoechst or DAPI.

### Annotation of EM data

Each EM volume (Nguyen et al., 2023; Wilson et al., 2019) was uploaded onto CATMAID (Saalfeld et al., 2009) server. Centrioles and cilia were manually located and annotated. Centrioles were marked at the center inside the distal end of the centriole and sphere with a 500 nm radius was placed at the node for 3D visualization. The initial node of the cilia skeleton was placed at the center of the base of the cilium (which is also the proximal end of the centriole). Subsequent cilia nodes tracked the center of the cilium, and a tag was added to the skeleton to mark the location where a pocket cilium exited the cell. Centriole vectors in the adult volumes originated at the center of the distal end of the cilium and terminated at the center of the proximal end (Sup. Figure 4B). The classifications used for annotating in the P7 and the adult volumes are listed in Supplementary Table 2. Although it is a rough approximation, in the P7 volume, layer boundaries do not correlate directly with cell depth because of the orientation of the sample during EM sectioning. Layer annotations for each cell were therefore manually determined by the immediate cell context. The code used to extract data from the CATMAID annotations can be found at: https://github.com/pattonw/centriole_data.

### Visualization and segmentation of EM data

Images and volumes were cropped from the EM volume in CATMAID. Images were rotated to similarly orient all centrioles. Where needed to improve image quality or to compensate for differences within the raw data, image brightness and contrast were adjusted using Fiji/ImageJ. For segmentation, image stacks were imported into Amira. Each structure was segmented in a separate layer before meshworks were generated and the images of the 3D structure were captured. Segmentation was approximated where image quality was uninterpretable.

### Graphing and statistics

All graphs were generated using Prism. Superplots were generated by overlaying average values from each animal into individual values, as explained in (Lord et al., 2020). Statistical significance was determined in Prism using ordinary one-way Anova using multiple comparison analysis with Turkey correction. Population mean was assessed at a 95% confidence interval and were considered significant at the following p values: 0.0.332 (*), 0.0021 (**), 00002 (***), <0.0001 (****).

---

**Supplemental Figure 1.**
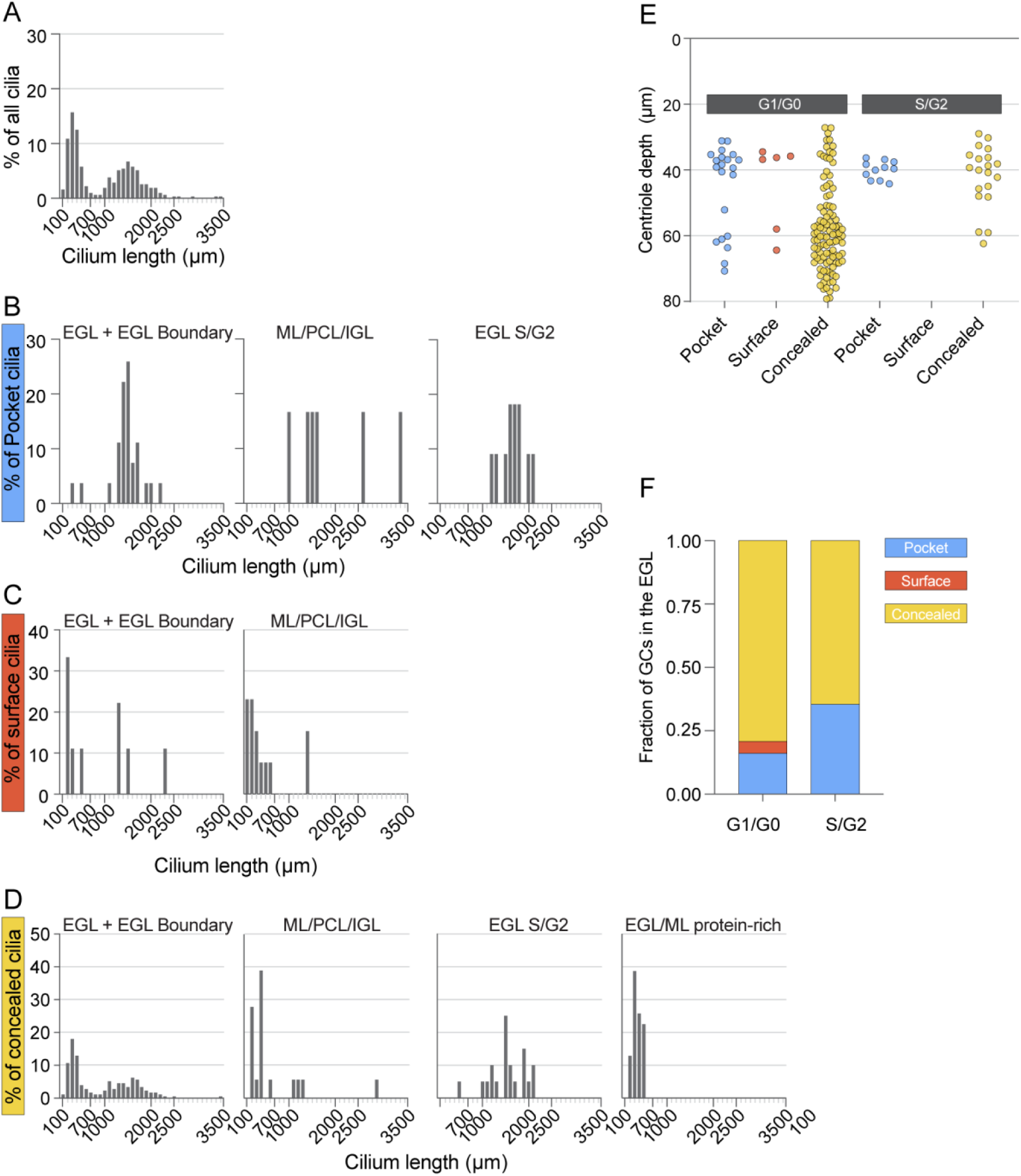
Bimodal distribution of cilia lengths. (A) Cilia from the P7 volume were binned by 100 nm and the distribution of cilia lengths was plotted as a percentile of total cilia. (B-D) The distribution of cilia lengths plotted in A is separated by pocket (B), surface (C), and concealed (D) cilia types. Within each cilium type the distributions are shown by cell layer with S/G2 cilia and protein-rich cilia plotted separately. There are no s/G2 cells with surface cilia and all protein-rich cilia are concealed. (E) The depth of the basal body for each cilium in the EGL is plotted based on cilium type. Cells in G1/G0 are on the left and cells in S/G2 are on the right. (F) The pocket, surface and concealed cilia in the EGL are plotted as a fraction of the total G1/G0 cells and S/G2 cells. Cells in the EGL boundary were excluded from E and F.

**Supplemental Figure 2.**
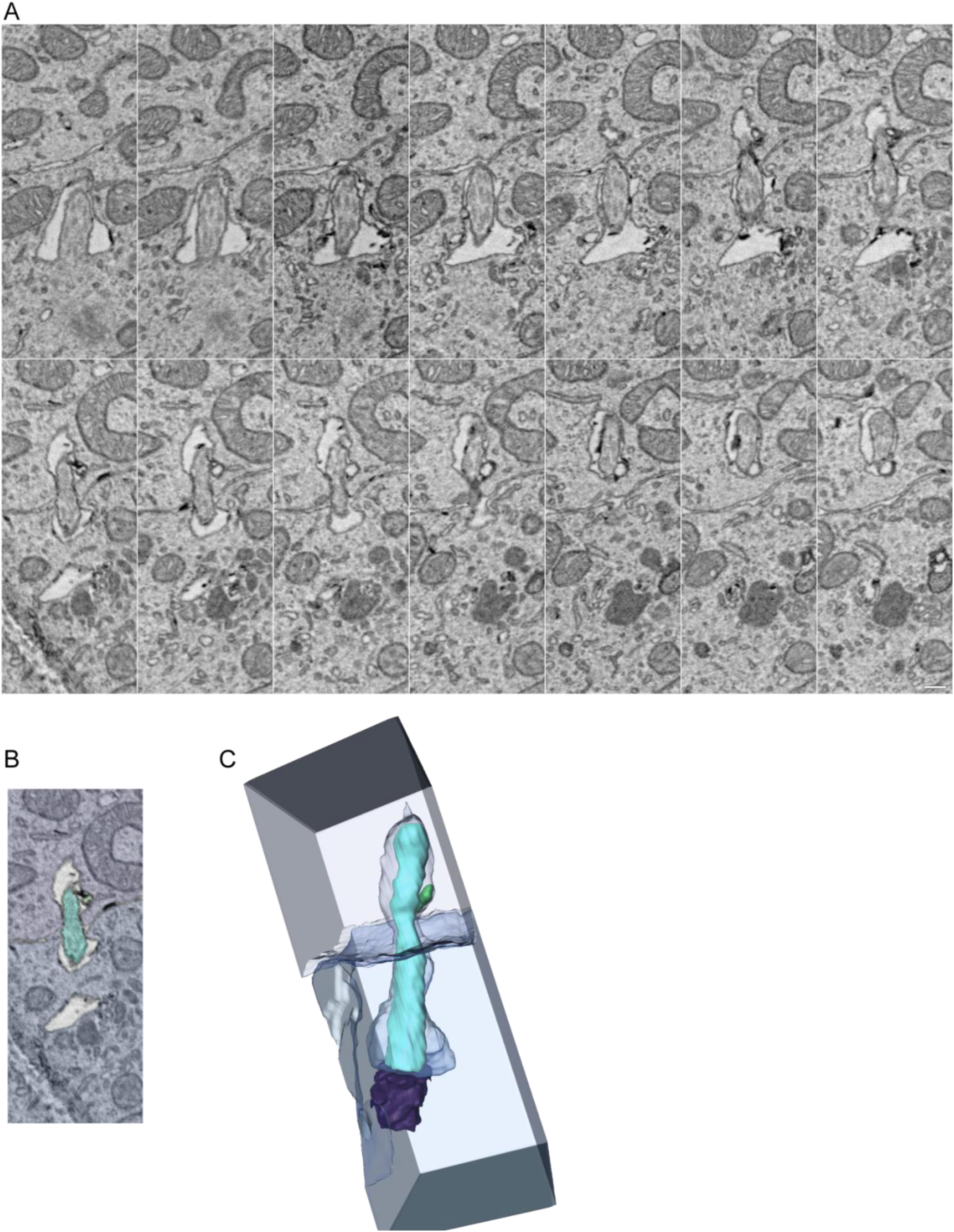
Cilium enveloped by an adjacent cell. (A) Serial EM sections of a pocket cilium that exits the cell in the bottom half of the image and is enveloped by the adjacent cell. Scale bar is 200 nm and z interval is 30 nm. (B) Single z slice from the sections in A colored to highlight the cilium (cyan), the cell of cilium origin (blue) and the enveloping cell (lavender). (C) 3D reconstruction of the same cilium. The basal body is purple and the membrane inclusion adjacent to the cilium is green.

**Supplemental Figure 3.**
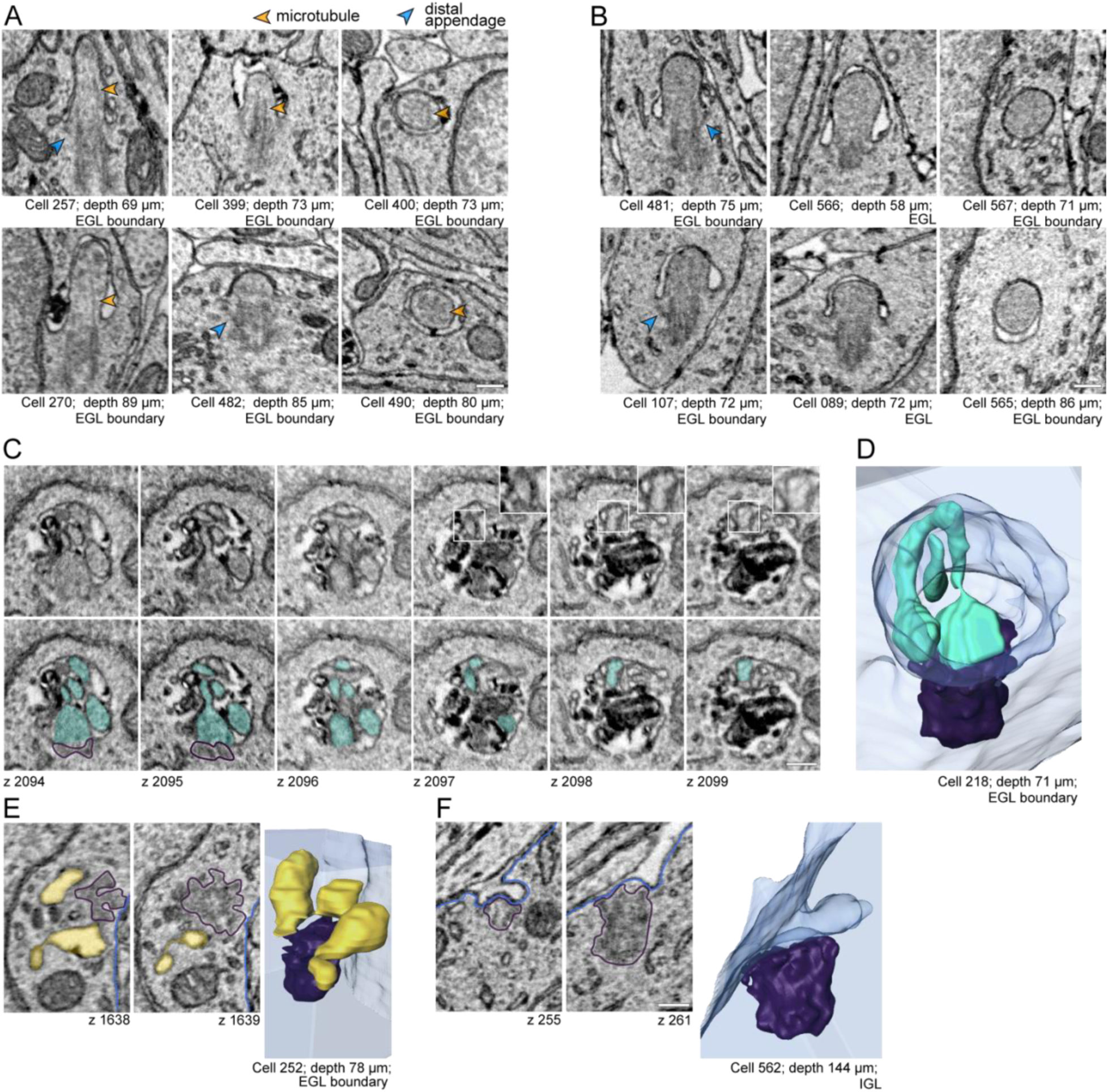
Cilium deconstruction and docking intermediates. (A-B) Representative images of short cilia with electron lucent (A) or electron rich (B) cilioplasm. Microtubules are highlighted with orange arrowheads and distal appendages with blue arrowheads. The scale bar is 200 nm. (C-D) A single cilium was observed with a constriction that could be indicative of cilium severing. Serial EM sections are shown in C. Microtubule singlets are visible in the insets on the top row and the cilium (and potential cilium fragment) are shaded cyan in the lower panel. A view of the 3D segmented image is presented in D. (E) A mother centriole in the EGL boundary that has three vesicles each docked at a different subset of distal appendages. (F) A centriole in the IGL with a membrane invagination anchored at distal appendages. Scale bars are 200 nm.

**Supplemental Figure 4.**
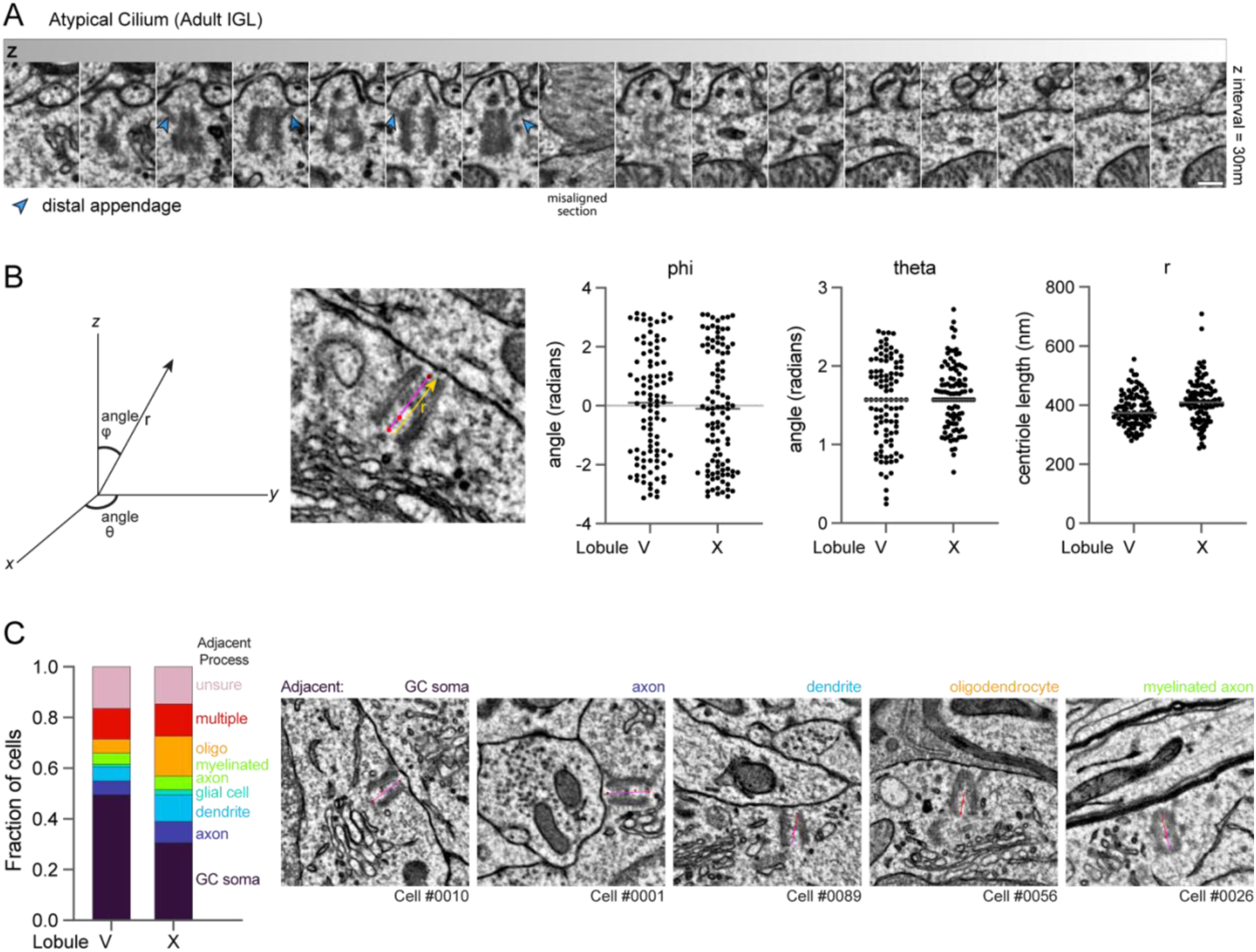
Docked centrioles in adult GCs lack directed orientation. (A) Serial EM sections display the only cilium annotated in the adult IGL volumes. Internal vesicles are present and the axoneme is not present or not resolved. Scale bar is 200 nm. (B) To assess polarity of centriole docking we generated a vector from proximal to distal within each mother centriole. We compared the vectors and found no bias in the orientation of docked centrioles. (C) The type of structure immediately adjacent to the plasma membrane where each GC mother centriole docked was determined. The distributions for the annotated centrioles in each adult dataset are plotted on the left and EM images on the right provide examples of centrioles docked adjacent to the indicated structures.

**Supplemental Figure 5.**
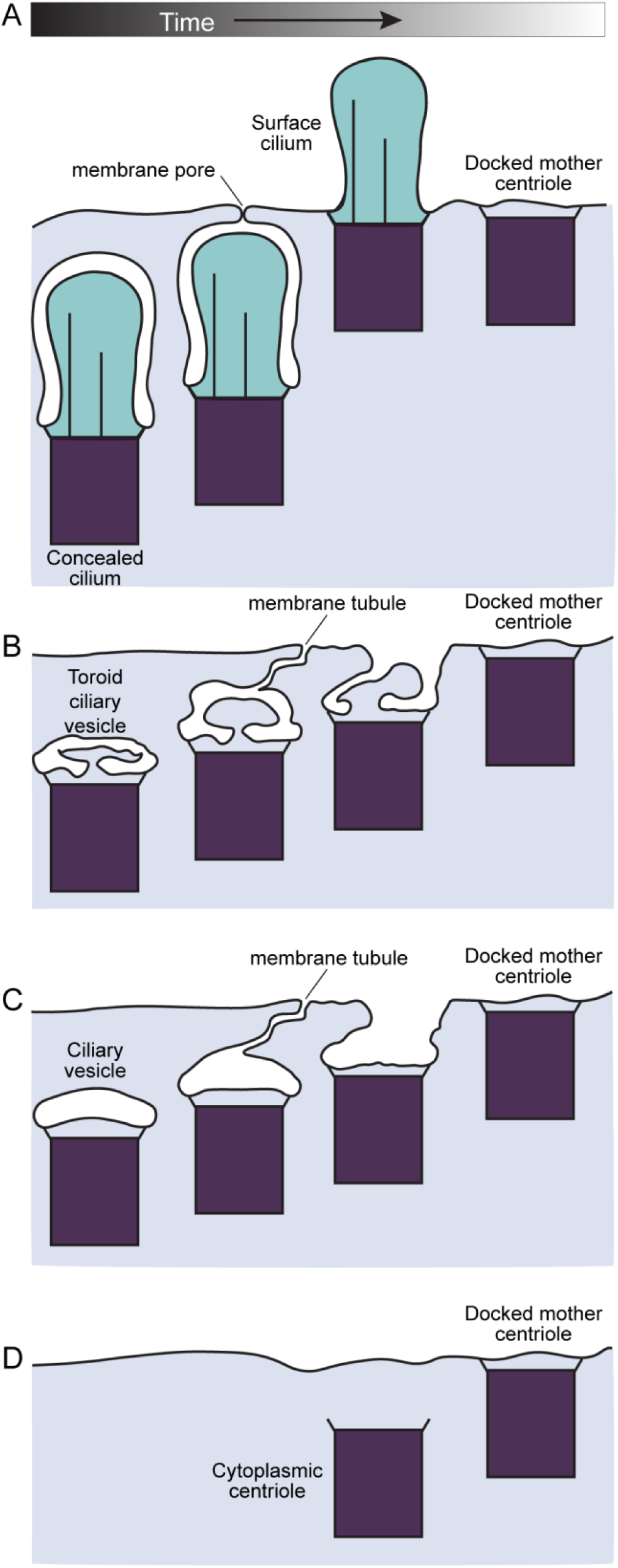
Centriole docking initiated at different stages in cilia deconstruction could account for the variety of intermediates observed. Differences between late-stage cilia/centrosome structures in differentiating cells suggest that instead of a linear deconstruction pathway, variance in the coordination of cilia deconstruction and mother centriole docking could generate multiple routes to mature, unciliated cells with docked mother centrioles. (A) Concealed cilia could access the plasma membrane through observed membrane pores. Opening of the pore before the cilium has been deconstructed could result in surface cilia, which could subsequently be completely disassembled. (B and C) Cilia deconstruction could proceed such that ciliary vesicles remain while centriole docking commences. Dynamic tubules could be key to unite the ciliary vesicle with the plasma membrane. (D) Conventional plasma membrane docking of cytoplasmic mother centrioles, which have completely deconstructed cilia, could proceed similar to centriole docking during surface cilia biogenesis and at the immune synapse.

**Supplemental Movie 1. Pocket Cilium in the EGL.**

This z series includes the entire pocket cilium of cell 451 shown in Figure 4A.

**Supplemental Movie 2. Surface Cilium in the IGL.**

This z series includes the entire surface cilium of cell 447 shown in Figure 4B.

**Supplemental Movie 3. Concealed Cilium in the EGL.**

This z series includes the entire concealed cilium of cell 518 shown in Figure 4C.

**Supplemental Movie 4. Concealed Cilium in the EGL boundary.**

This z series includes the entire pocket cilium of cell 368 shown in Figure 4D.

**Supplemental Movie 5. Centriole with ciliary vesicle in EGL boundary.**

This z series includes the entire mother centriole of cell 564 shown in Figure 5F.

**Supplemental Movie 6. Centriole with toroid-shaped ciliary vesicle in EGL.**

This z series includes the entire mother centriole of cell 232 shown in Figure 5G.

**Supplemental Movie 7. Centriole with ciliary vesicle in IGL.**

This z series includes the entire mother centriole of cell 436 shown in Figure 5H.

**Supplemental Movie 8. Docked centriole with membrane invaginations in ML.**

This z series includes the entire mother centriole of cell 021 shown in Figure 5I.

**Supplemental Movie 9. Docked centriole in IGL.**

This z series includes the entire mother centriole of cell 056 shown in Figure 5J.

**Supplemental Movie 10. Tethered centriole in IGL.**

This z series includes the entire mother centriole of cell 049 shown in Figure 5K.

**Supplemental Movie 11. Cytoplasmic centriole in IGL.**

This z series includes the entire mother centriole of cell 064 shown in Figure 5L.

